# Identification of Potential DNA Gyrase Inhibitors: Virtual Screening, Extra-Precision Docking and Molecular Dynamics Simulation Study

**DOI:** 10.1101/2022.11.06.515362

**Authors:** Avinash Kumar, Chakrawarti Prasun, Ekta Rathi, Maya S. Nair, Suvarna G Kini

## Abstract

DNA gyrase brings negative supercoils into DNA and loosens up certain positive supercoils that collect during replication and transcription and is a notable antibacterial target. To fight against the menace of antibiotic-resistant bacterial infections, we have employed various computational tools like high throughput virtual screening (HTVS), standard precision (SP), extra precision (XP), and molecular dynamics (MD) simulation studies to identify some potential DNA gyrase inhibitors. A focused library of 5968 anti-bacterial compounds was screened using the HTVS docking protocol of the glide module of Maestro. The top 200 docked compounds were further filtered using SP and XP docking protocols and their free binding energies were calculated using MM-GBSA studies. Binding and stability of the top two compounds which showed better docking score than the co-crystallized ligand (clorobiocin) of DNA gyrase (PDB ID: 1KZN) was further probed by MD simulation of 100 ns using GROMACS. MD simulation study suggested that the compounds AM1 and AM5 form a stable complex with DNA gyrase with a good number of hydrogen bonds. XP docking study showed that interaction with the crucial amino acids for compounds AM1 and AM5 was like the co-crystallized ligand. These compounds were also predicted to be drug-like molecules with good water solubility and excellent absorption profiles. Based on the above studies, herein we report compounds AM1 (1R,3S)-1-(2-((3-(ammoniomethyl)phenyl)amino)-2-oxoethyl)-3-carbamoylpiperidin-1-ium and AM5 (1’S,2s,4R)-4-ammonio-6-ethyl-1’-methylspiro[chromane-2,4’-piperidin]-1’-ium as potential DNA gyrase inhibitors which can be further developed as a potential drug against the menace of antibiotic resistance.

## 1. INTRODUCTION

DNA gyrase is a topoisomerase II, an ATP-dependent enzyme that is essential for the processes of chromosome segregation, transcription, and replication. It belongs to gyrase, heat-shock protein 90 (Hsp 90) histidine kinase MutL (GHKL), protein kinases and the DNA mismatch repair protein MutL (Mismatch from replication recognized by mutL)) family of enzymes (Kirchhausen, Wang, & Harrison, 1985). It is one of the most extensively researched and verified targets for the creation of novel antibacterial therapies. This enzyme is a good target for the development of antibacterial therapeutics with selective toxicity due to its lack in the mammalian organism and its critical function in the bacterial DNA replication cycle (Khan et al., 2018). Gyrase A and Gyrase B are the two subunits that make up the catalytically active heterotetrameric enzyme (i.e. A2B2). While the B subunit (DNA gyrase B) has ATPase activity and supplies enough energy for the DNA supercoiling, the A subunit is responsible for breaking and rejoining the double DNA strand (Champoux, 2003; Dutta & Inouye, 2000).

Given that DNA gyrase is a highly desirable pharmacological target, many inhibitors have been created and characterised, with quinolones and aminocoumarins being the most extensively researched substances (Maxwell, 1997). Presently, the only DNA gyrase inhibitors utilised in clinical practice belong to the class of substances known as 6-fluoroquinolones. In-depth research in this area has been sparked by the variety of adverse effects and developing bacterial resistance in the absence of new antibacterial agents. By screening chemical libraries, new chemical entities with various scaffolds and DNA gyrase inhibitory capabilities have been identified. These compounds could make promising leads for the development of antibacterial agents (Khan et al., 2018).

Recently, Pakamwong *et al*., have identified 2,2’-((2-hydroxy-3-(5-methyl-2,3-diphenyl-1H-indol-1-yl)propyl)azanediyl)bis(ethan-1-ol) as potential DNA gyrase inhibitor having an antimycobacterial activity (Pakamwong et al., 2022). Govender *et al*., have reported some potent spiropyrimidinetrione derivatives as DNA gyrase inhibitors with potent activity against mycobacterium tuberculosis (Govender et al., 2022). Alfonso *et al*., have discovered a few novels and structurally diverse DNA gyrase inhibitors through a fluorescence-based high-throughput screening assay (Alfonso et al., 2022). Saleh *et al*., have employed experimental and computational (molecular docking) studies to discover some cyclic diphenyl phosphonates as DNA gyrase inhibitors for fluoroquinolone-resistant pathogens (Saleh et al., 2022).

Recently, several authors have employed computational tools to identify novel inhibitors of various targets (Kumar, Rai, Rathi, Agarwal, & Kini, 2020; Meena et al., 2021; Mukherjee et al., 2022; Rathi, Kumar, & Kini, 2020). Jakhar *et al*., have employed quantitative structure-activity relationship (QSAR) modelling to identify a few novel DNA gyrase inhibitors (Jakhar, Khichi, Kumar, Dangi, & Kumar Chhillar, 2022). Hasan *et al*., have carried out *in silico* analysis of ciprofloxacin derivatives as potential DNA gyrase inhibitors (Hasan et al., 2021). Mohammed *et al*., have synthesized and performed docking studies of a few novel N4-piperazinyl ciprofloxacin analogues as potential DNA gyrase inhibitors (Mohammed et al., 2022). We wanted to identify some novel chemotypes which already might have antimicrobial activity, and be developed as DNA gyrase inhibitors. Therefore, in the present study, we have employed various computational tools like high throughput virtual screening (HTVS), molecular docking, and molecular dynamics (MD) simulation studies to identify some potential DNA gyrase inhibitors. The findings of the current study might serve as a hit that can be optimized for DNA gyrase inhibition and in turn developed as a potent antimicrobial agent against antibiotic-resistant strains.

## 2. Materials and Methods

All molecular modelling experiments and DFT studies were carried out using Maestro (v11.4) on an HP desktop with Linux Ubuntu 18.04.1 LTS as the operating system. Maestro is a small-molecule drug discovery suite from Schrodinger that has several integrated molecular modelling tools like Glide, Phase, Desmond, QikProp, LigPrep etc. MD simulation was carried out using GROMACS 2018.3 on the LINUX system with NVIDIA GPU support. DFT studies were calculated at the B3LYP level of DFT using a 6-31+G(d,p) basis set in the gas phase with no restrictions in the Gaussian 03 package of programs.

### 2.1 Protein Preparation

The protein structure (PDB ID 1KZN) was imported from the protein databank using the import option in the protein preparation wizard tool of the Maestro interface(Berman, Henrick, & Nakamura, 2003; Lafitte et al., 2002). The protein preparation wizard tool has a three-step integrated process where the imported protein structures are first pre-processed to fill hydrogens or any missing residues using an integrated Phase module. In the preprocessing step, bond orders are also corrected and then water molecules beyond 5Å were removed, added hydrogens were optimized and finally, protein structure was minimized using the OPLS3e (optimized potentials for liquid simulations) force field(Madhavi Sastry, Adzhigirey, Day, Annabhimoju, & Sherman, 2013).

### 2.2 Database Preparation

For the present study, a focused library of 5968 anti-bacterial compounds was downloaded from the Asinex database(“Asinex.com - Antibacterial - Research Areas - Screening Libraries,” n.d.). The Asinex database provides a variety of libraries of compounds for screening purposes. All the compounds were downloaded in .mol2 format and imported using the LigPrep tool of Maestro(Madhavi Sastry et al., 2013). Lowest energy 3D structures were generated for all the imported structures with corrected chiralities using the LigPrep tool. This exercise was carried out using an OPLS3e forcefield (Roos et al., 2019).

### 2.3 High Throughput Virtual Screening

HTVS mode in the Glide module of Maestro was employed to carry out high throughput virtual screening. HTVS mode is the fastest docking mode in the Glide panel. It uses very less computational power as its scoring function does not use the explicit-water technology and descriptors as used in XP mode. Hence, HTVS is preferred for the screening of a large library of compounds. But its algorithm is not able to weed out false positives. HTVS docking is a grid-based docking system and hence, a receptor grid was first generated for the minimized protein structure (1KZN). The grid was generated using the co-crystallized ligand with the protein as a reference ligand(Friesner et al., 2004).

### 2.4 Molecular Docking

Based on the docking score, the compounds were ranked in the HTVS experiment and the top 200 compounds were taken up for docking in the standard precision (SP) mode of Glide. The SP mode uses a sampling method based on anchor and refined growth. The top 50 compounds based on the docking score in SP docking were selected for XP (extra precision) docking. XP mode is less forgiving than SP mode and penalties are levied for violations like inadequate solvation as predicted by statistical results(Richard A. Friesner et al., 2006).

### 2.5 Free Binding Energy Calculations

The top five compounds from XP docking studies were put for free binding energy calculations using the MM-GBSA(Molecular Mechanics, the Generalized Born model and Solvent Accessibility) tool of the Prime module of Maestro. Prime utilizes the VSGB 2.0 solvation model and OPLS3e force field to simulate these interactions. Ligands were ranked based on MM-GBSA ΔG bind scores (Genheden & Ryde, 2015).

### 2.6 MD Simulation Study

The top two compounds based on the above studies were taken up for MD simulation for more insight into the interaction and stability of the protein-ligand complex.GROMACS (version 2018.3) was used with AMBER ff99SB-ILDN forcefield and tip3p solvent model (Berendsen, van der Spoel, & van Drunen, 1995). Antechamber tool of AMBERTOOLS was employed for generating the ligand topology. The system was neutralized by adding counter ions (Na^+^/Cl). The solvated complex was subjected to energy minimization by employing conjugate gradient methods and the steepest descent (SD) algorithm.Post minimization, two-phase equilibration was carried out and the equilibrated complex was put for MD simulation at 300K and 1 bar for 100 ns with a time step of 2 fs.

### 2.7 ADME Analysis

ADME (absorption, distribution, metabolism, and excretion) analysis was done using SwissADME which is a web-based tool. SwissADME helps us to forecast ADME parameters, pharmacokinetic qualities, druglike nature and medicinal chemistry friendliness by harnessing its ability to compute physicochemical descriptors(Daina, Michielin, & Zoete, 2017).

## 3. Results and Discussion

### 3.1 Molecular Docking and MM-GBSA Studies

From a library of 5968 compounds, initial screening was done using HTVS docking and based on the docking score, the top 200 compounds were selected for SP docking studies. As SP docking algorithms are more refined than HTVS docking, therefore the second level of screening was done using SP mode. The top 50 compounds based on the docking score in the SP mode, were selected for docking in the XP mode. The docking results were compared with the standard ligand (clorobiocin) co-crystallized with the target protein. The docking score of clorobiocin was found to be −6.078 kcal/mol and it showed H-bond interactions with GLY77 and THR165 amino acid residues (Figure 1f). Hydrophobic interactions were observed with VAL43, ALA47, VAL71, ILE78, PRO79, ILE90, ALA96, VAL118, GLY119, VAL120, MET166, AND VAL167 residues. Polar interactions were observed with ASN46, GLN72, HIS95, and THR165residues. Charged (negative interactions were observed with ASP49, GLU50, and ASP73 amino acid residues.

**Figure 1.**
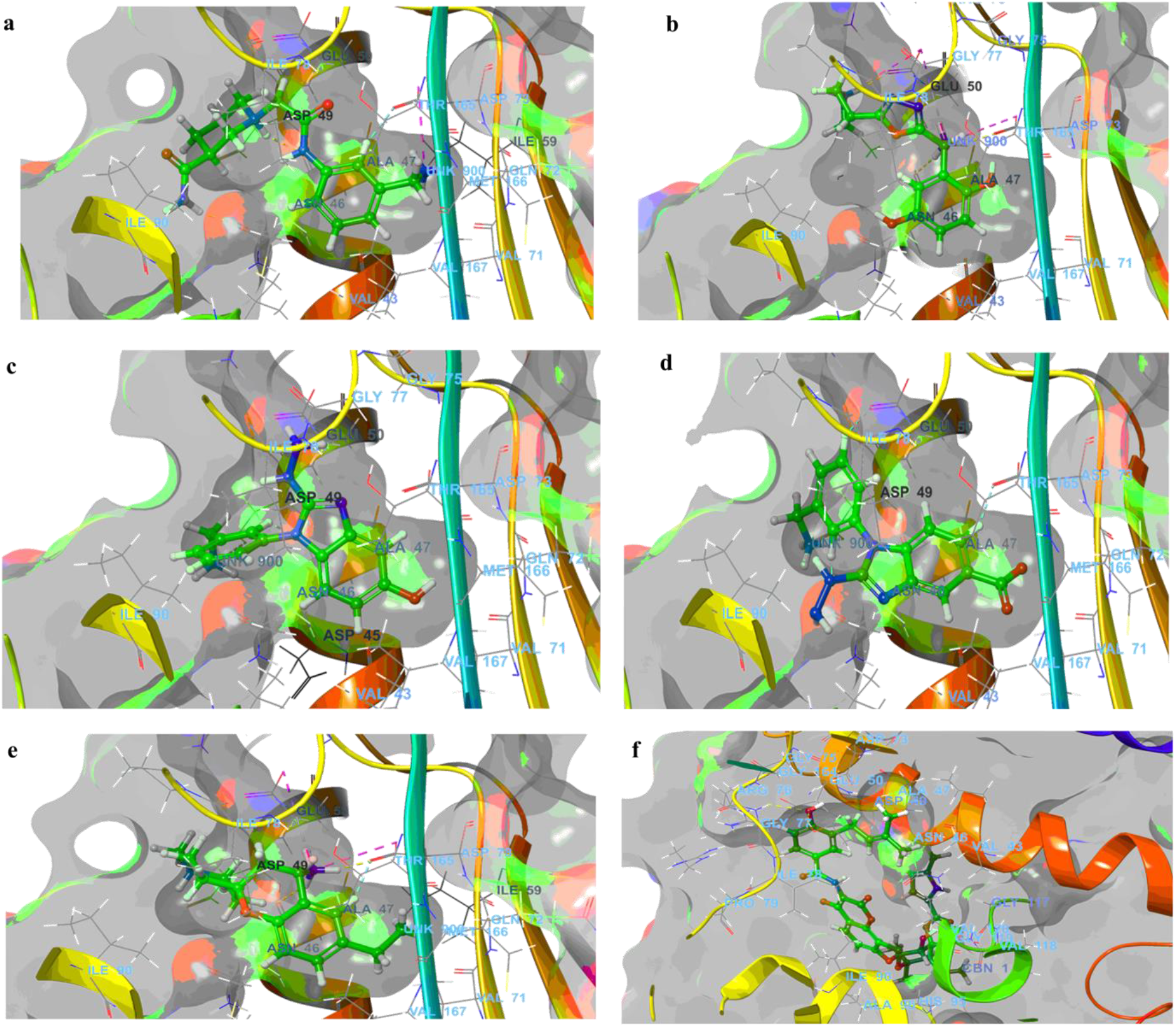
Ligand interaction diagram of top five docked compounds (in XP mode) and clorobiocin (co-crystallized ligand). (a) AM1 (b) AM2 (c) AM3 (d) AM4 (e) AM5 (f) clorobiocin

In comparison to clorobiocin, all five selected compounds showed a better docking score which was >-8.0 kcal/mol (Table 1). Compound AM5 showed the best docking score of −10.160kcal/mol while compound AM4 showed the least docking score of −8.007 kcal/mol which was still better than clorobiocin. Non-bonding interactions like H-bonding, hydrophobic, polar interactions etc. were also similar to clorobiocin as shown in Table 1 and Figure 1. Salt-bridge type interactions were also observed with all of these ligands except AM3 and AM4 with ASP73 residue which was not observed for clorobiocin. Compounds AM3 and AM4 showed salt-bridge interaction with ASP49 residues. The ligand interaction diagram of all the top ligands has been shown in Figure 1.

**Table 1.**
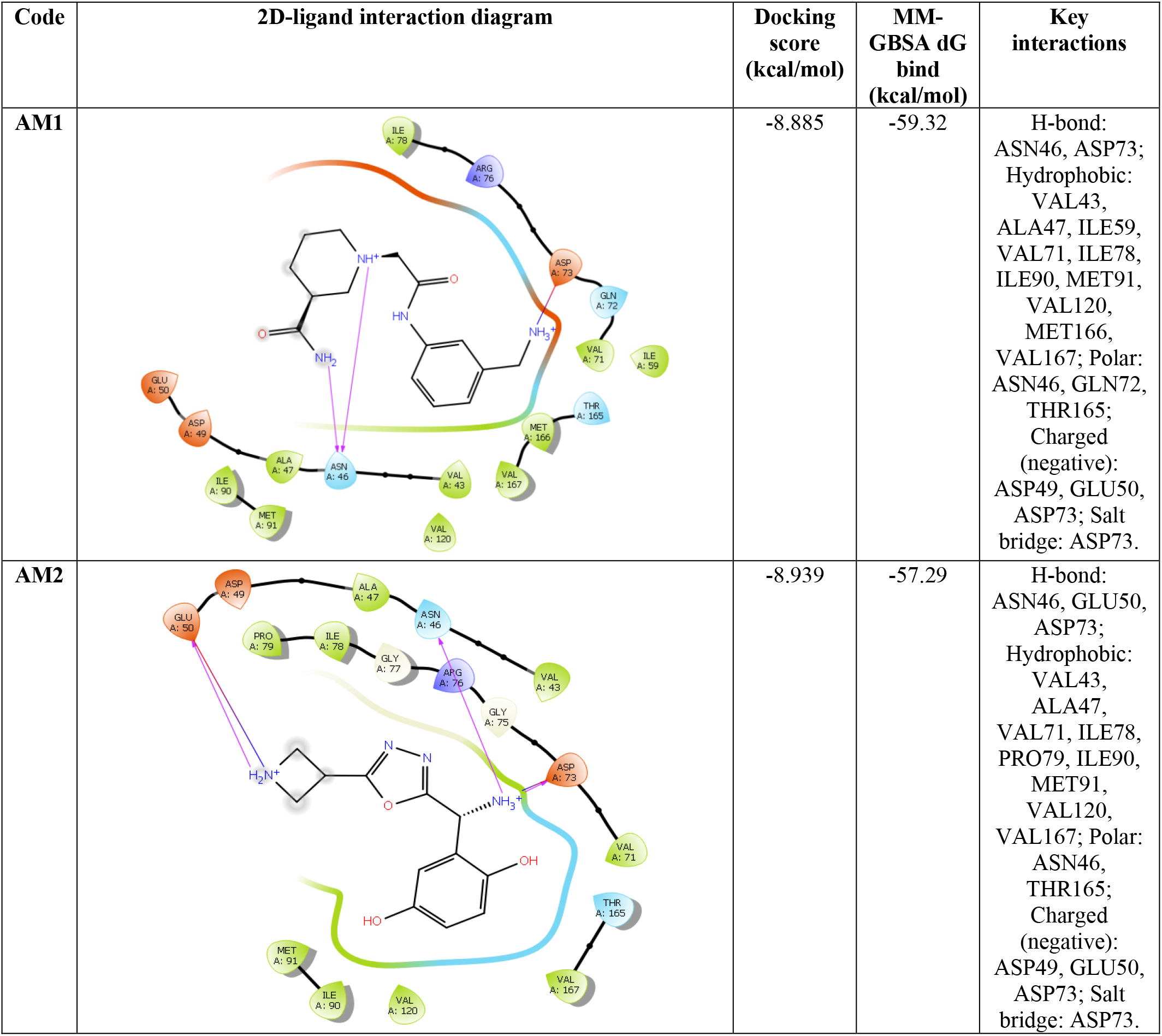
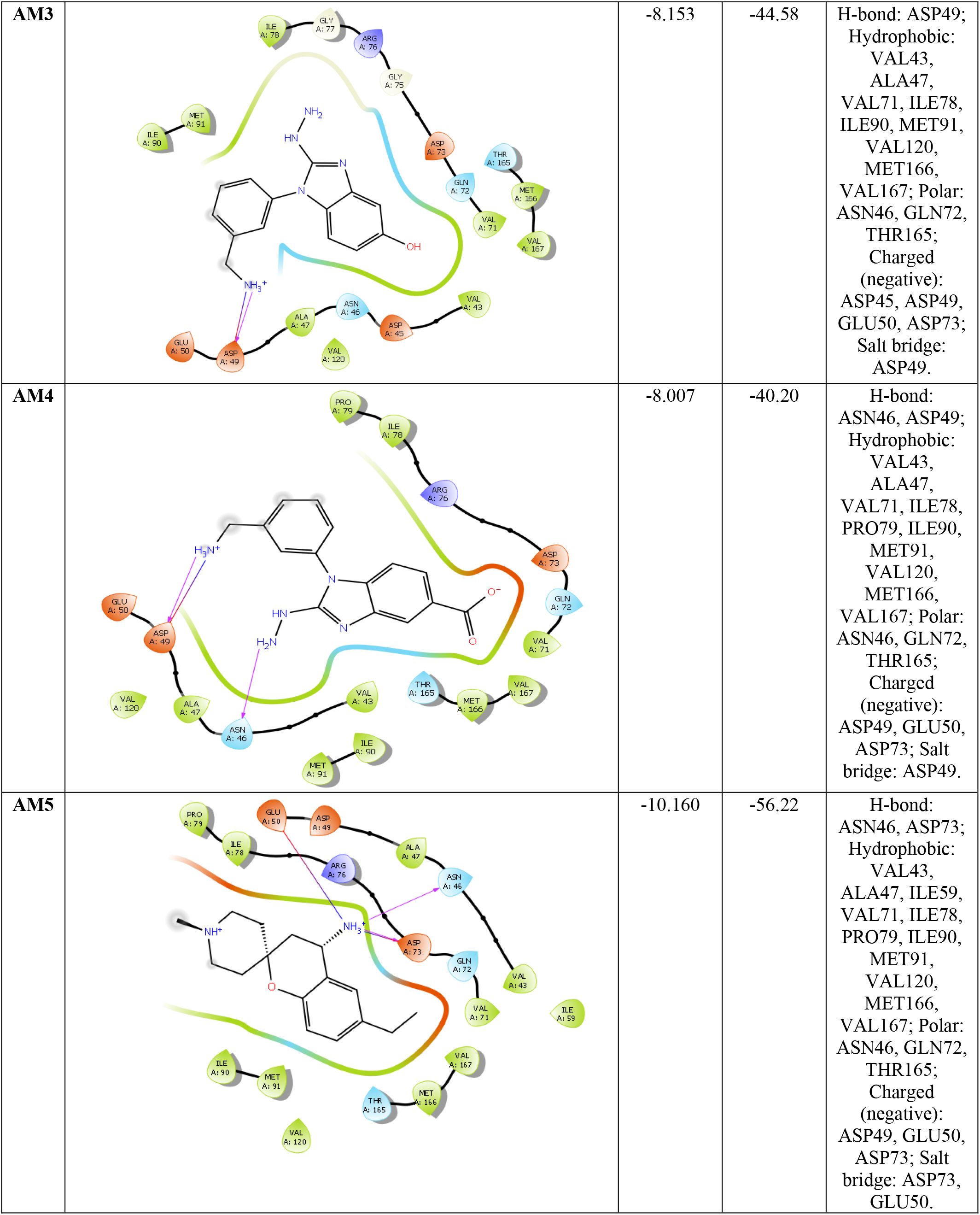
Docking results (in XP mode) of the top five selected ligands

| Code | 2D-ligand interaction diagram | Docking score (kcal/mol) | MM-GBSA dG bind (kcal/mol) | Key interactions |
| --- | --- | --- | --- | --- |
| AM1 |  | -8.885 | -59.32 | H-bond:<br>ASN46, ASP73;<br>Hydrophobic:<br>VAL43,<br>ALA47, ILE59,<br>VAL71, ILE78,<br>ILE90, MET91,<br>VAL120,<br>MET166,<br>VAL167; Polar:<br>ASN46, GLN72,<br>THR165;<br>Charged<br>(negative):<br>ASP49, GLU50,<br>ASP73; Salt<br>bridge: ASP73. |
| AM2 |  | -8.939 | -57.29 | H-bond:<br>ASN46, GLU50,<br>ASP73;<br>Hydrophobic:<br>VAL43,<br>ALA47,<br>VAL71, ILE78,<br>PRO79, ILE90,<br>MET91,<br>VAL120,<br>VAL167; Polar:<br>ASN46,<br>THR165;<br>Charged<br>(negative):<br>ASP49, GLU50,<br>ASP73; Salt<br>bridge: ASP73. |
| AM3 |  | -8.153 | -44.58 | H-bond: ASP49;<br>Hydrophobic:<br>VAL43,<br>ALA47,<br>VAL71, ILE78,<br>ILE90, MET91,<br>VAL120,<br>MET166,<br>VAL167; Polar:<br>ASN46, GLN72,<br>THR165;<br>Charged<br>(negative):<br>ASP45, ASP49,<br>GLU50, ASP73;<br>Salt bridge:<br>ASP49. |
| AM4 |  | -8.007 | -40.20 | H-bond:<br>ASN46, ASP49;<br>Hydrophobic:<br>VAL43,<br>ALA47,<br>VAL71, ILE78,<br>PRO79, ILE90,<br>MET91,<br>VAL120,<br>MET166,<br>VAL167; Polar:<br>ASN46, GLN72,<br>THR165;<br>Charged<br>(negative):<br>ASP49, GLU50,<br>ASP73; Salt<br>bridge: ASP49. |
| AM5 |  | -10.160 | -56.22 | H-bond:<br>ASN46, ASP73;<br>Hydrophobic:<br>VAL43,<br>ALA47, ILE59,<br>VAL71, ILE78,<br>PRO79, ILE90,<br>MET91,<br>VAL120,<br>MET166,<br>VAL167; Polar:<br>ASN46, GLN72,<br>THR165;<br>Charged<br>(negative):<br>ASP49, GLU50,<br>ASP73; Salt<br>bridge: ASP73,<br>GLU50. |

Free binding energy calculations were also carried out for the top five ligands in the complex with the protein. The compounds were ranked based on dG bind energy and all the selected compounds showed favourable binding free energy profiles (Table 1). Compound AM1 showed the best dG bind energy of −59.32 kcal/mol while compound AM4 showed the lowest value of −40.20 kcal/mol. Clorobiocin showed a dG bind score of −64.36 kcal/mol and hence, compounds AM1, AM2 and AM5 were found to show a comparable free binding energy profile with the standard Clorobicin.

### 3.2 MD Simulation Study

The stability of the DNA gyrase and its identified inhibitors (AM1 and AM5) based on the docking studies was further validated through an MD simulation study for 100 ns. Docking studies are devoid of any water molecule and there is no control over the temperature and pressure parameters also. Therefore, the MD simulation study gives a better understanding of the complex stability in the aqueous environment and at room temperature and pressure. Various information like the conformational changes in the ligands, protein residues and changes in their interaction during the simulation period, stability etc. are collected during the simulation and are represented as various plots like RMSD (root mean square deviation), SASA (solvent accessible surface area), H-bond plot, RMSF (root mean square fluctuation), radius of gyration etc.

#### 3.2.1 RMSD Analysis

RMSD plots are very crucial in MD simulation studies. It gives an insight into the overall stability of the protein-ligand complex which can be analyzed over two phases i.e., the equilibrium phase and the production phase. The first frame is taken as the reference frame and all subsequent frames are superimposed over it to obtain the RMSD plot to time. As shown in figure 2, the RMSD of Cα- atoms were computed at the beginning and for the AM1-DNA gyrase complex (black colour in figure 2), the system reached the equilibrium phase at around 10 ns. For the AM5-DNA gyrase complex (red colour in figure 2), equilibrium was attained at around 20ns. During the post-equilibrium phase, both the complexes remained stable throughout the simulation period as shown by the average RMSD deviation between 0.1 to 0.4 nm. The RMSD plot suggested that the compounds AM1 and AM5 might form a stable complex with DNA gyrase.

**Figure 2.**
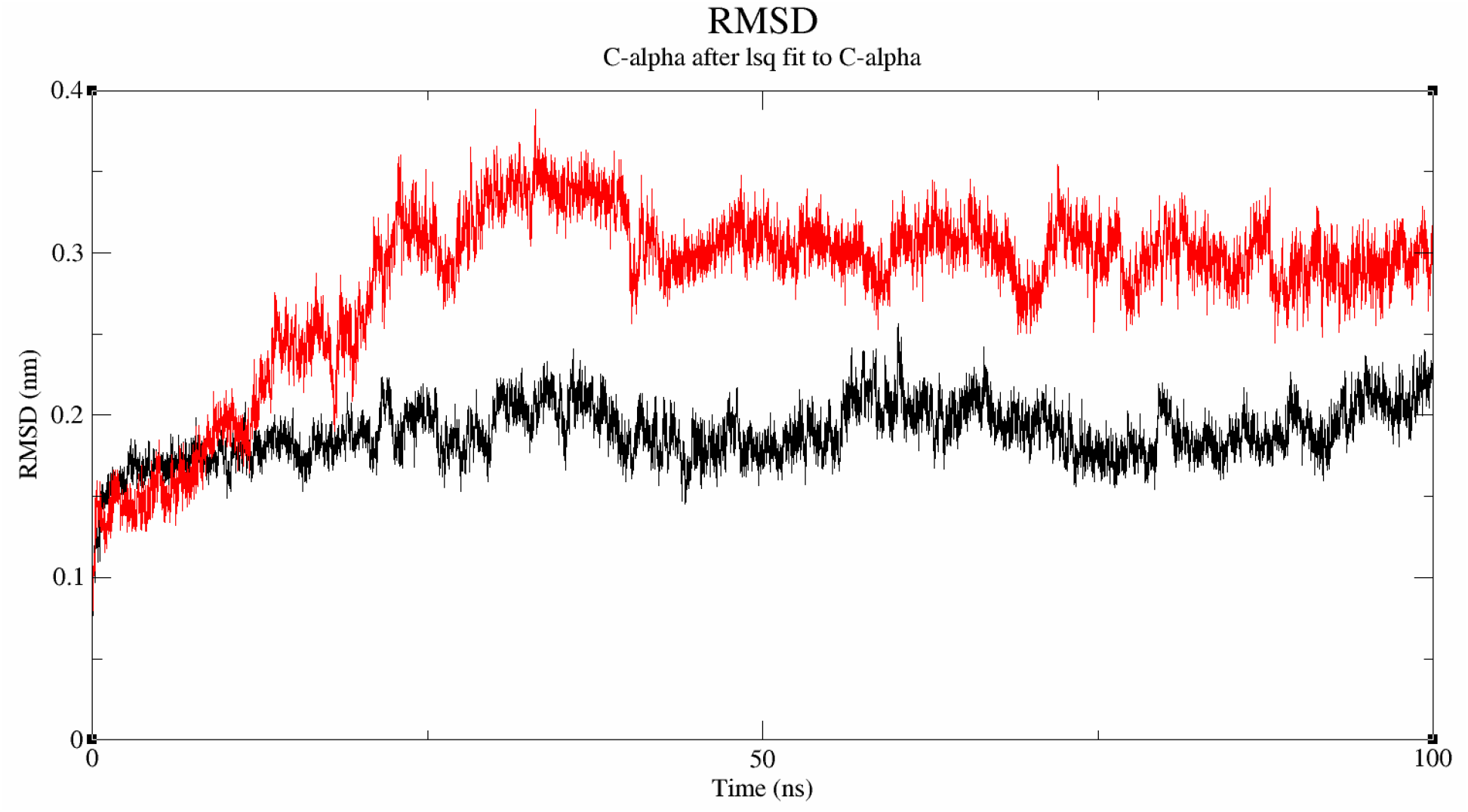
RMSD plot of DNA gyrase and ligand complex. The black colour shows the AM1-DNA gyrase complex and the red colour shows the AM5-DNA gyrase complex.

#### 3.2.2 SASA and H-bond Analysis

Intermolecular hydrogen bonding is one of the most important interactions between protein and ligand and plays a key role in the stability of the complex. The distribution and number of H-bonding between the ligand and protein during MD simulation are depicted in Figure 3a. For the AM1-DNA gyrase complex (black colour), there were a few poses which showed 7 H-bonds while many of the poses showed 5-6 H-bonds. The average density of the H-bond was strongly distributed around 3 and 4 which suggests that AM1-DNA gyrase might form a stable complex. For the AM5-DNA gyrase complex, a few poses showed 3 to 4 H-bonds while the average H-bonding interaction was 2 implying that the AM5-DNA gyrase complex might also be stable through the simulation period.

**Figure 3.**
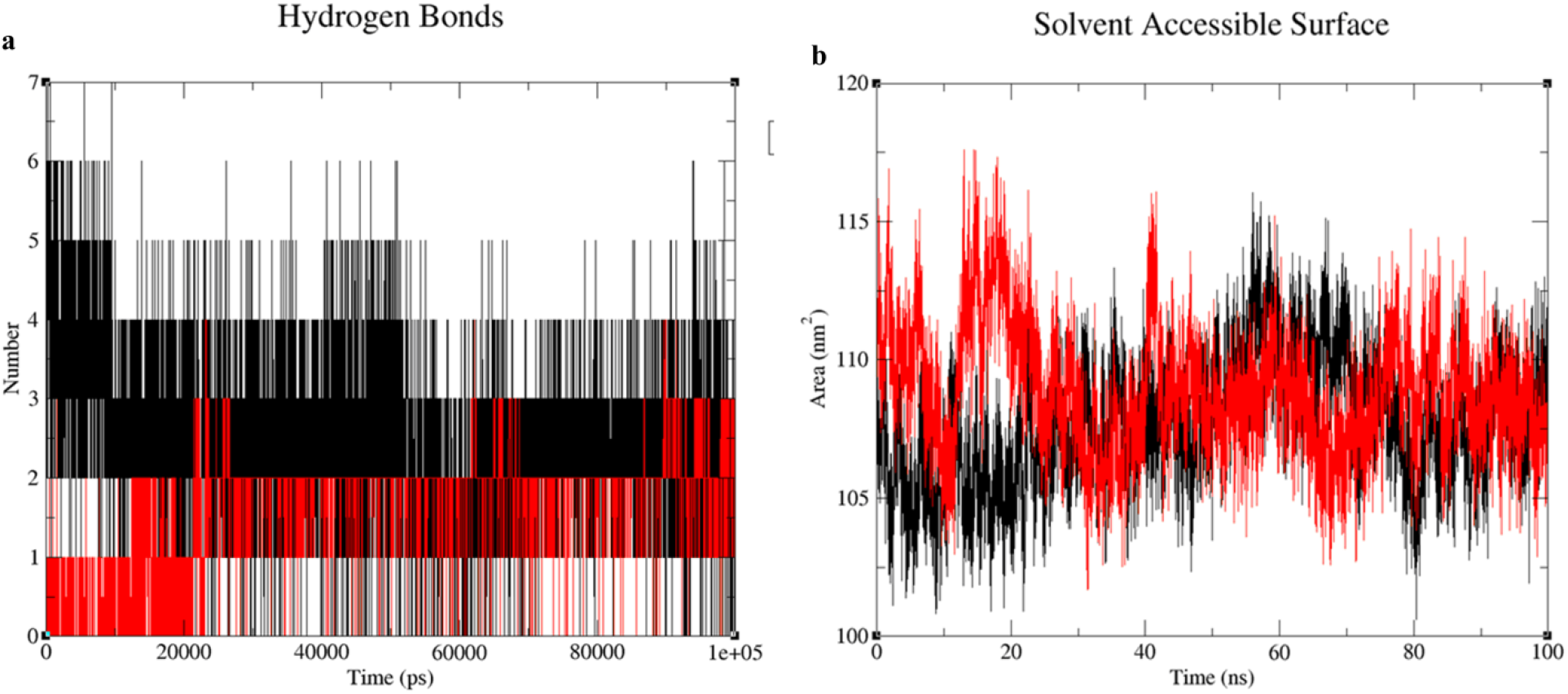
**(a)** H-bond plot of AM1-DNA gyrase complex (black) and AM5-DNA gyrase complex (red). **(b)** SASA plot of AM1-DNA gyrase complex (black) and AM5-DNA gyrase complex (red).

SASA plot represents the change inaccessibility of protein to the solvent. It gives an insight into the analysis of the folding pattern of the inner hydrophobic core to the outer hydrophilic surface of the protein structure. As depicted in Figure 3b, the SASA for AM1-DNA gyrase complex (black) and AM1-DNA gyrase complex (red) was in the range of 100-120 nm^2^ which suggests that there were no major changes in the folding pattern of the protein due to binding with the ligands and hence, might have remained stable throughout the simulation period.

### 3.3 ADME Analysis

Initially, we thought to do ADME profiling of the compounds to select only drug-like molecules from the library of 5968 compounds. But then we didn’t want to miss out on any potent inhibitor of DNA gyrase because of the unfavourable ADME profile. As this is an *in-silico* study, so the potent DNA gyrase inhibitors can be identified at this stage and then they can be optimized for their pharmacokinetic properties. Therefore, we ran ADME profiling of only the top five compounds that were identified using XP docking studies. Several parameters were computed for all the compounds and some of them like molecular weight, the number of H-bond donors and acceptors, polar surface area (PSA), solubility, gastrointestinal (GI) absorption, blood-brain-barrier (BBB) permeation, Lipinski’s rule of five violations, PAINS alert, lead-likeness, and synthetic feasibility have been reported in Table 2. All the compounds were lead-like molecules with no violation of the rule of five. There were no PAINS alerts for any compounds except AM2. They also showed good GI absorption (especially AM3, AM4, and AM5). All the compounds were predicted to be water-soluble and none of them could cross BBB. They were also predicted to be Pgp substrates except for AM2 and AM4 and none of them inhibited CYP1A2 and its other isoforms like CYP2C19, CYP2C9, CYP2D6, and CYP3A4. Synthetic accessibility scores also suggested that these compounds can be synthesized. Based on the ADME predictions, the selected compounds AM1 and AM5 seem to be drug-like inhibitors of the DNA gyrase enzyme.

**Table 2.** ADME profile of the top five selected compounds as predicted by SwissADME.

| Code | MW | HbD | HbA | TPSA | ESOL Class | GI absorption | BBB permeant | P-gp substrate | CYP1A2 inhibitor | Rule of Five | PAINS alert | Lead-likeness | Synthetic Accessibility |
| --- | --- | --- | --- | --- | --- | --- | --- | --- | --- | --- | --- | --- | --- |
| AM1 | 292.38 | 4 | 2 | 104.27 | Very Soluble | Low | No | Yes | No | 0 | 0 | Yes | 2.73 |
| AM2 | 264.28 | 4 | 5 | 123.63 | Very Soluble | Low | No | No | No | 0 | 1 | Yes | 3.23 |
| AM3 | 270.31 | 4 | 3 | 103.74 | Very Soluble | High | No | Yes | No | 0 | 0 | Yes | 2.41 |
| AM4 | 297.31 | 3 | 4 | 123.64 | Very Soluble | High | No | No | No | 0 | 0 | Yes | 2.45 |
| AM5 | 262.39 | 2 | 1 | 41.31 | Soluble | High | No | Yes | No | 0 | 0 | Yes | 4.03 |

MW: molecular weight; HbD: hydrogen bond donor; HbA: hydrogen bond acceptor; TPSA: topological polar surface area; P-gp: P-glycoprotein.

## 4. Conclusion

In the present study, we have employed HTVS docking to screen a focused library of antibacterial compounds to identify potential DNA gyrase inhibitors. The top 200 compounds were further filtered with the help of more advanced docking algorithms like SP and XP docking. Finally, based on the docking score and MM-GBSA studies, the top two compounds were selected for MD simulation studies. The docking score of the co-crystallized ligand (clorobiocin) was found to be −6.078 kcal/mol and in its comparison, the top five selected compounds showed a docking score of >-8.0 kcal/mol. MD simulation studies also suggested that the ligand-receptor complex between compounds AM1, AM5, and DNA gyrase would be stable as supported by the RMSD and H-bond plots. ADME studies also reinforced the findings that compounds AM1 and AM5 have an acceptable pharmacokinetic profile and are drug-like molecules. Based on the above studies, herein we report compounds AM1 (1R,3S)-1-(2-((3-(ammoniomethyl)phenyl)amino)-2-oxoethyl)-3-carbamoylpiperidin-1-ium and AM5 (1’S,2s,4R)-4-ammonio-6-ethyl-1’-methylspiro[chromane-2,4’-piperidin]-1’-ium as potential DNA gyrase inhibitors which can be further developed as a potential drug against the menace of antibiotic resistance.

## Conflict of Interest

The authors declare no conflict of interest.

## ACKNOWLEDGMENTS

The authors are also grateful to Manipal-Schrodinger Centre for Molecular Simulations, Manipal College of Pharmaceutical Sciences, MAHE, Manipal for providing facilities to carry out molecular modelling studies.

## Notes

### Competing Interest Statement

The authors have declared no competing interest.

